# Non-paradoxical evolutionary stability of the recombination initiation landscape in Saccharomycetes

**DOI:** 10.1101/023176

**Authors:** Isabel Lam, Scott Keeney

## Abstract

The nonrandom distribution of meiotic recombination shapes heredity and genetic diversification. A widely held view is that individual hotspots — favored sites of recombination initiation — are always ephemeral because they evolve rapidly toward extinction. An alternative view, often ignored or dismissed as implausible, predicts conservation of the positions of hotspots if they are chromosomal features under selective constraint, such as gene promoters. Here we empirically test opposite predictions of these theories by comparing genome-wide maps of meiotic recombination initiation from widely divergent species in the *Saccharomyces* clade. We find that the frequent overlap of hotspots with promoters is true of the species tested and, consequently, hotspot positions are well conserved. Remarkably, however, the relative strength of individual hotspots is also highly conserved, as are larger-scale features of the distribution of recombination initiation. This stability, not predicted by prior models, suggests that the particular shape of the yeast recombination landscape is adaptive, and helps in understanding evolutionary dynamics of recombination in other species.

## One Sentence Summary

Hotspots for meiotic double-strand breaks decay rapidly in some species but are maintained over long evolutionary times in yeast, where they are targeted to functional genomic features.

## Main Text

DNA double-strand breaks (DSBs) generated by Spo11 protein initiate meiotic recombination, which in turn alters genetic linkage and promotes pairing and accurate segregation of homologous chromosomes (*1*). DSBs are not distributed randomly, occurring often within narrow (typically < 200 bp wide in budding yeast) regions called hotspots (*2*). Theoretical work exploring evolutionary dynamics has led to a prevailing hypothesis, the “hotspot paradox”, that predicts rapid hotspot extinction (*3-8*). This view rests on biased gene conversion: the broken chromosome copies genetic information from its uncut homolog (Fig. 1A). As a result, hotspot alleles with different DSB activity deviate from a Mendelian segregation ratio, with colder (less active) alleles overrepresented in offspring. Such meiotic drive is observed in yeast (*9*) and humans (*10*) and predicts that mutations that reduce or eliminate hotspot activity will be rapidly fixed in populations, while hotspot-activating mutations should be rapidly extinguished (*3, 5, 11*). The paradox is that hotspots for DSBs and recombination exist at all.

**Fig. 1.**
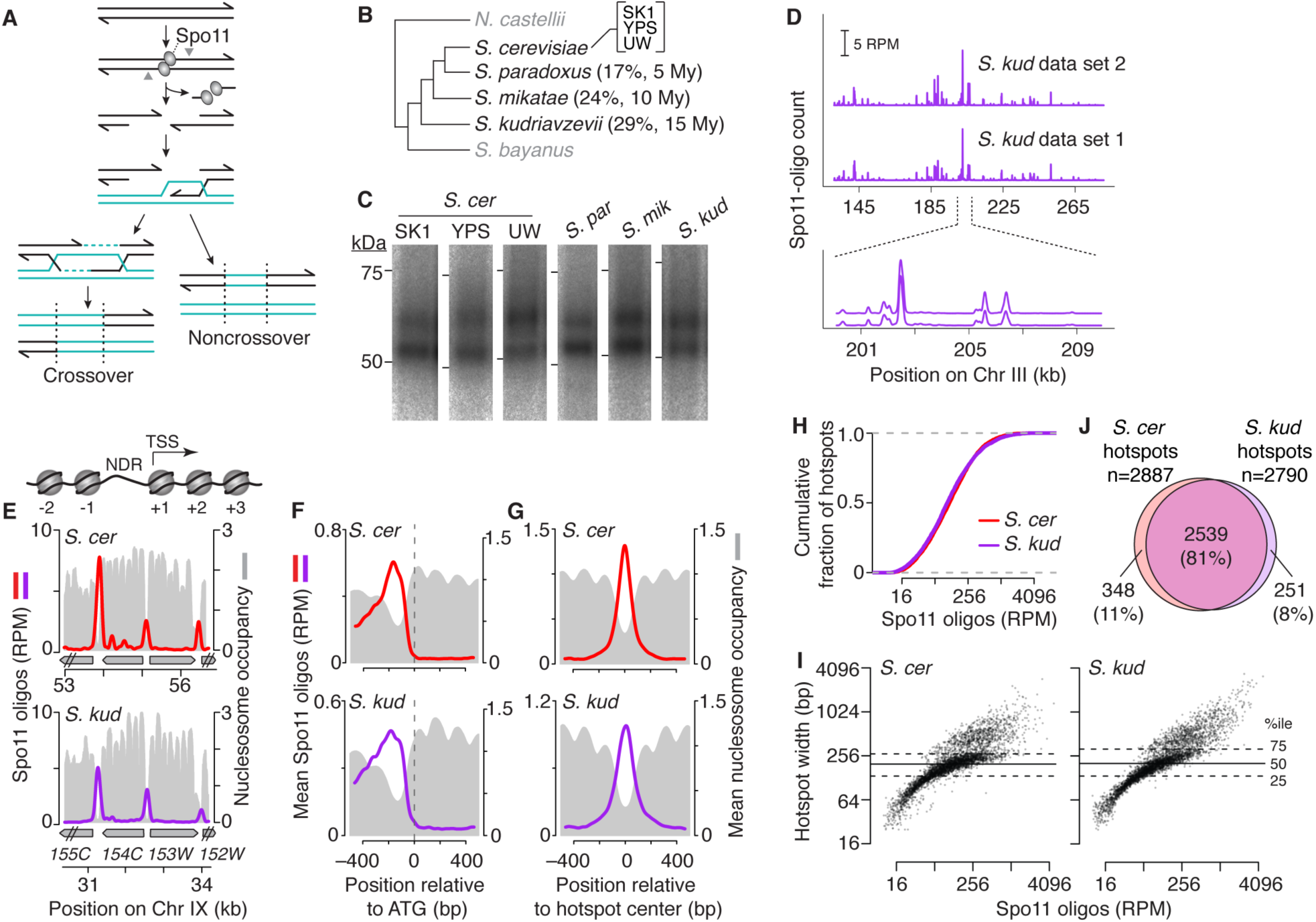
Conserved targeting of DSBs to promoters. (A) Meiotic recombination. Spo11 generates covalent protein-linked DSBs; endonucleolytic cleavage (grey arrowheads) liberates Spo11 bound to short oligos exploited to map DSBs. DSB ends are resected and repaired to yield crossover or non-crossover products. The broken chromosome (black) copies information from the uncut allele (teal). (B) *Saccharomyces* phylogeny (schematic of outgroup (*N. castelli*)-rooted neighbor-joining tree using Rad51 protein alignment). Black, species/strains in this study; genic percent sequence divergence from *S. cerevisiae* (*49*) and estimated time since last common ancestor (18, 50) are shown. YPS, YPS128 strain; UW, UWOPS03-461.4 strain. (C) Conserved sizes of Spo11-oligo complexes. Autoradiographs of immunoprecipitated, radiolabelled complexes separated by SDS-PAGE. (D) Reproducibility of *S. kudriavzevii* Spo11-oligo maps. RPM, reads per million mapped. (E) Overlap of DSB hotspots with promoter NDRs is evolutionarily conserved. The cartoon depicts typical yeast promoter chromatin structure, with an NDR upstream of the transcription start site (TSS). Sample region (around *YIL154C*) compares Spo11 oligos with the nucleosome map (MNase-seq read depth relative to genome average). (F) Average Spo11-oligo and nucleosome profiles around start codons (*S. cer* SK1, n=5766 genes; *S. kud*, n=5578). (G) Average Spo11-oligo and nucleosome profiles at hotspots (*S. cer* SK1, n=4099; *S. kud*, n=3976). (H) Hotspot intensity varies over similar smooth continua in *S. cerevisiae* (SK1) and *S. kudriavzevii*. (I) Similar distributions of widths vs. Spo11-oligo counts in hotspots from *S. cerevisiae* (SK1) and *S. kudriavzevii*. (J) Conservation of hotspot position. Most promoter-containing IGRs hosting Spo11-oligo hotspots in *S. cerevisiae* (SK1) also had hotspots in *S. kudriavzevii* (see **fig. S3G** for more detail and other species). Spo11-oligo profiles were smoothed with 201-bp (panels D,E) or 75-bp (panels F,G) Hann window.

An alternative view predicts that hotspot positions can be preserved if Spo11 targets genomic features that are under selective constraint to maintain functions unrelated to their roles as hotspots (9, 12, 13). This hypothesis derives from the fact that most DSB hotspots in the budding yeast *S. cerevisiae* correspond to promoter-containing intergenic regions (13). However, theoretical studies have considered this implausible as a mechanism to preserve hotspots (3, 11). Instead, many studies take as their starting point the assumption that hotspot lifespan must always be short and that the fine-scale recombination initiation landscape will always be highly dynamic over long evolutionary times (4, 6-8, 14). This assumption is appropriate for humans and most other mammals (discussed further below), but has not been evaluated for other taxa.

To distinguish between these models, we asked whether the DSB landscape is conserved in yeast. A prior attempt to address this issue used population genetic data to deduce a recombination map in *S. paradoxus* and compared it to *S. cerevisiae* (15). Partial conservation was inferred, but the data had insufficient resolution and sensitivity to detect individual hotspots and were confined to one small chromosome that may not be representative of most of the yeast genome. We overcame these limitations by comparing high-resolution, whole-genome DSB maps between more widely diverged *Saccharomyces* species, as well as between *S. cerevisiae* strains from different lineages (the laboratory strain SK1 and wild-derived strains YPS128 and UWOPS03-461.4) (16) (Fig. 1B, **Table S1**). DSB maps were generated by deep-sequencing of DNA oligonucleotides (oligos) that remain covalently bound to Spo11 as a byproduct of DSB formation and processing (13, 17) (Fig. 1A, **Table S2**).

The *Saccharomyces sensu stricto* clade shared a last common ancestor ∼20 million years ago (18). We chose species at varying evolutionary distances from *S. cerevisiae* (Fig. 1B) ranging from *S. paradoxus*, with protein coding sequence divergence from *S. cerevisiae* comparable to that between humans and mouse, to *S. kudriavzevii*, roughly as distant as mammals from birds (19). The *S. cerevisiae* strains chosen display 0.5–0.7% sequence divergence, comparable to the polymorphism density between humans and chimpanzees. Most differences are simple sequence polymorphisms, with few large-scale structural differences aside from one discussed below (20). The *Saccharomyces* karyotype is thus an evolutionarily stable, successful one.

These yeasts all underwent synchronous and efficient meiosis (**fig. S1A**), hence SK1 is not anomalous in this regard. As in *S. cerevisiae*, two major size classes of Spo11-oligo complexes were observed (Fig. 1C, fig. S1B), reflecting oligos of similar length distributions (**fig. S1C**). Each oligo is a tag recording precisely where Spo11 generated a DSB, and maps based on deep sequencing Spo11 oligos agree spatially and quantitatively with direct detection of DSBs by Southern blot (13, 21, 22). Biological replicate maps were highly reproducible (Fig. 1D, fig. S2) and most sequenced reads (>98%) mapped uniquely to their respective genomes (**Table S2**).

We first asked whether targeting of promoters is conserved. To aid analysis, we generated nucleosome occupancy maps by sequencing micrococcal nuclease-resistant DNA (MNase-seq) from meiotic cultures. In *S. cerevisiae*, DSBs form preferentially in promoter-associated nucleosome-depleted regions (NDRs) (13, 23), and promoter chromatin structure is strongly conserved in other *Saccharomyces* species (24-26). Spo11 oligos were highly enriched in promoter NDRs in all species tested, whether examined at individual locations (Fig. 1E, fig. S3A), or averaged across annotated genes (Fig. 1F, fig. S3B). Many Spo11 oligos mapped to promoter-containing intergenic regions (IGRs) (i.e., those flanked by divergent or tandemly oriented transcription units), whereas few mapped to convergent IGRs (i.e., those lacking promoters) or within genes (**fig. S3D**). Clearly, Spo11 preference for promoters is a stable feature of the *Saccharomyces* DSB landscape.

Similar numbers of Spo11-oligo hotspots (∼4000) were identified in all species (**Table S3**). When ranked by heat (Spo11-oligo count), hotspots formed a smooth continuum over a wide range in both *S. cerevisiae* and *S. kudriavzevii* (Fig. 1H), with nearly superimposable cumulative curves in all species (**fig. S3E**). Hence, the distribution of DSBs among hotspots is the same. Hotspots decisively tended toward low nucleosome occupancy (Fig. 1G, fig. S3C) consistent with open chromatin structure providing a window of opportunity for Spo11 to access DNA (27). The distribution of hotspot widths was also nearly identical, with wider hotspots tending to have more Spo11 oligos (Fig. 1I, fig. S3F). Conserved hotspot width agrees with previously established conservation of NDR width (24-26). Most important, the majority of hotspots overlapped the same promoter-containing IGRs in all species examined (Fig. 1J, fig. S3G). The low frequency of sex and outcrossing in yeasts could slow hotspot extinction compared to obligately outcrossed species (15, 28), but the yeasts examined here have had ample sexual generations to allow biased gene conversion to erode hotspots (estimated >200,000 outcrossed sexual generations since divergence of *S. cerevisiae* from *S. kudriavzevii*, comparable to the number of human sexual generations since divergence from chimpanzees; see Methods). Thus, as predicted (9), DSB hotspot locations can be preserved when the underlying chromosome architecture being targeted is conserved.

We next examined conservation of hotspot heat. The hotspot paradox predicts that hotspot activity will be in a state of flux: even if hotspot locations are conserved, they should vary widely in absolute and relative strength. Furthermore, the rapidity of hotspot extinction should scale with hotspot heat, because alleles that experience high DSB frequency provide more chances for loss for a given number of sexual generations (3-5, 11, 29). The selective constraint model is agnostic in this regard. If cis-acting sequence polymorphisms can quantitatively modulate DSB formation without ablating Spo11 targeting per se (experimentally shown to be a plausible proposition (e.g., 30)), then hotspot heats will be free to change and will do so rapidly (13). On the other hand, if DSB frequency (not just position) is tied to selectively constrained aspects of the targeted feature, or if DSB frequency is itself constrained, then hotspot heats will tend to be conserved.

To address this question, we summed Spo11 oligos mapping within 3426 promoter-containing IGRs that could be stringently and unambiguously matched between species based on conservation of the flanking coding sequences (Fig. 2A, Table S4). (This accounts for 81% of divergent and tandem IGRs and 83% of promoter-proximal DSB hotspots in *S. cerevisiae*, so most of the relevant Spo11-targeted genomic space is included. An IGR-centric approach is preferable to relying on more arbitrary hotspot definitions; see Methods.) Intra-IGR Spo11-oligo counts were highly similar between *S. cerevisiae* strains, with correlation coefficients (0.89– 0.92) nearly as high as for comparisons between biological replicates (0.97–1.00) (Fig. 2B,C, fig. S4A, and data not shown). Thus, yeast intra-species variation in local DSB heat is low despite ∼0.7–1% median sequence divergence within these IGRs. Strong positive correlations were also found between species, with little change in correlation strength over large evolutionary distances (Fig. 2B,C, fig. S4A). Moreover, the hottest 1% of promoter IGRs in *S. cerevisiae* SK1 (based on Spo11-oligo count) were highly enriched among the hottest IGRs in other strains and species, with a median percentile ranking within the top 5% even in *S. kudriavzevii* (Fig. 2D,E). This was only modestly greater than the extent of conservation of the coldest IGRs (Fig. 2D), whereas theoretical modeling of the effects that biased gene conversion alone should have on hotspot longevity predicts that strong hotspots are much less likely to be shared between species than weak ones (5). We did find specific examples where strong hotspots in one species were substantially weaker in other species (Fig. 2F), so there is no absolute barrier to evolutionary changes in DSB activity. But the behavior of most IGRs leads us to conclude that the hottest hotspots present in the last *Saccharomyces* common ancestor have tended to retain their places at the top of the list of Spo11 targets, and that it has been rare for ancestrally cold promoter regions to acquire strong hotspot activity.

**Fig. 2.**
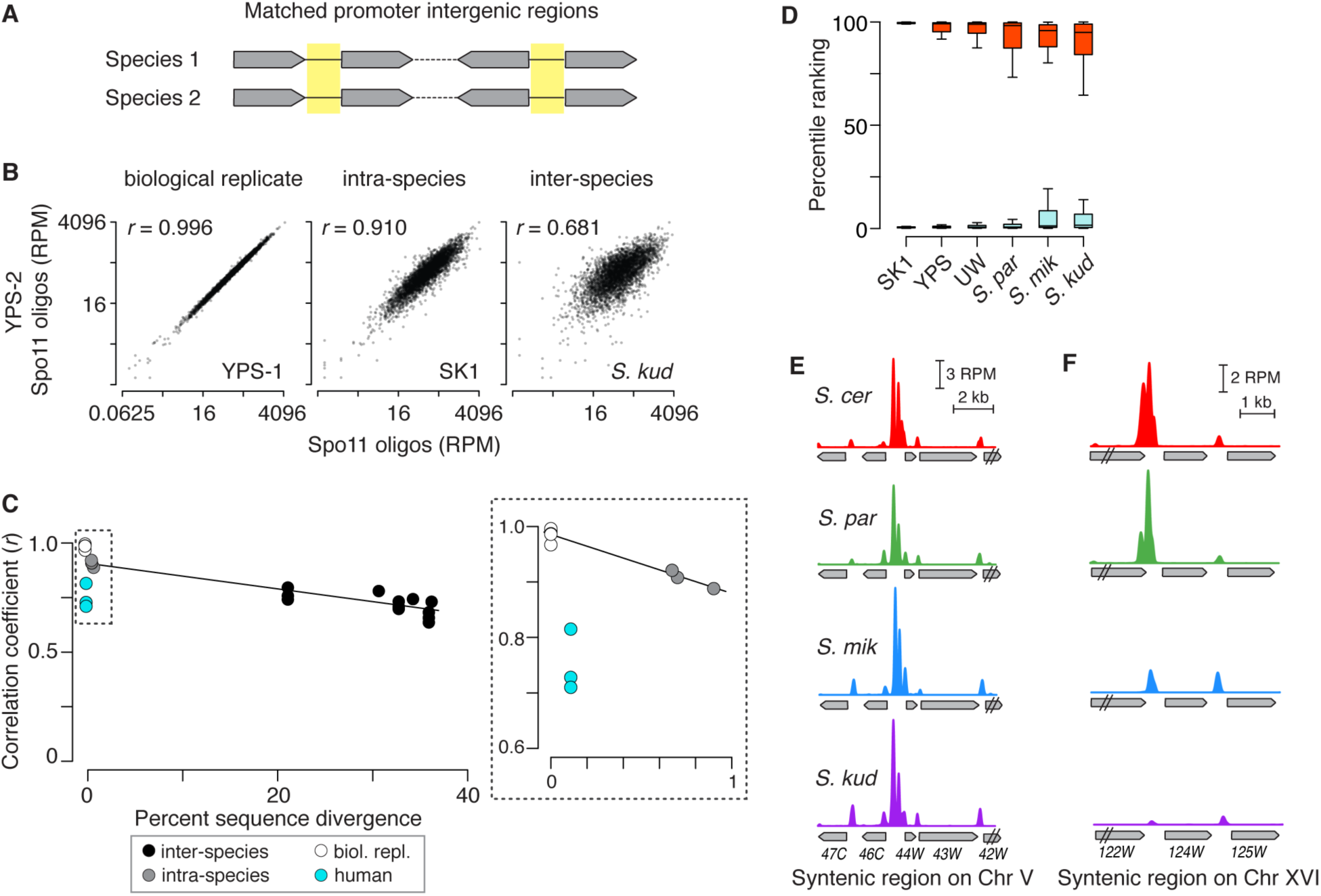
Conservation of hotspot strength. (A) Promoter-containing IGRs were matched between species based on conservation of both flanking genes. (B) Comparison of Spo11-oligo counts within 3426 IGRs that were matched in all four species. Correlation coefficients for the log_2_- transformed data are shown (Pearson’s *r*). (C) Spo11-oligo counts in promoter IGRs remain highly correlated despite wide sequence divergence. Correlation coefficients (as in panel B) are plotted against the median sequence divergence within the IGRs, which is substantially greater than the coding sequence divergence (Fig. 1B). Black line highlights relation between the yeast comparisons; it is not a regression line. Human data (from ref. 34) are for three men with identical or similar *PRDM9* alleles (37,345 hotspots (see **fig. S4B**); each had ∼0.1% sequence difference from the reference genome (34)). (D) The hottest hotspots have stayed hot, and the coldest have stayed cold. Percentile rankings in other strains and species are shown for the matched promoter IGRs with the most (red) and least (cyan) Spo11 oligos in SK1 (top and bottom 1%). Box plots are as in **fig. S3D**. (E,F) Examples of a strong Spo11-oligo hotspot from SK1 whose heat is conserved (E, *YEL046C*) and one whose heat is not (F, *YPR124W*).

This high degree of hotspot conservation differs markedly from the situation in most mammals, where PRDM9 provides a different solution to the hotspot paradox (31). PRDM9 is a histone methyltransferase with an array of Zn-finger modules that rapidly evolve new DNA binding specificity. The DNA motifs it recognizes, which have no known intrinsic function, are lost quickly from genomes of humans and mice because of meiotic drive from biased gene conversion (29, 32, 33), but periodic appearance of new *PRDM9* alleles with different sequence specificity creates new hotspots suddenly and thus redraws the recombination initiation landscape (31). Accordingly, DSB hotspot heat between men sharing the same or similar *PRDM9* alleles (34) was less conserved than between *S. cerevisiae* strains despite much greater sequence identity (Fig. 2C, fig. S4B). This difference is consistent with PRDM9 motif erosion contributing to variation in hotspot strength between individuals (34).

Hotspots are only one level of non-randomness in the DSB landscape in that they reside within larger domains of intrinsically greater or lesser DSB potential (12, 13). Conservation has been noted for the distribution of crossover recombination over broad regions in several taxa (14), but DSB distributions have not been evaluated. We therefore asked if large-scale features of the DSB landscape are also conserved in yeast. Spo11-oligo maps demonstrated that DSB suppression near telomeres and centromeres (13, 27) is preserved across *Saccharomyces* evolution (Fig. 3A,B, fig. S5A,B). This behavior is not surprising, as recombination in these subchromosomal domains can interfere with genome integrity (subtelomeric regions are rife with repetitive DNA elements that can undergo nonallelic homologous recombination (35), and crossing over too close to centromeres can provoke meiotic segregation errors (36)). More remarkably, however, Spo11-oligo counts were highly correlated when compared in ∼20-kb segments within larger blocks of synteny across the remaining interstitial portions of chromosomes (Fig. 3C,D, fig. S5C-F, Table S5). Thus, the domain structure of the DSB landscape is also evolutionarily stable. As in *S. cerevisiae*, Spo11-oligo counts correlated positively with G+C content of DNA in each species tested, with similar scale-dependent patterns of weaker correlation at short distances (∼1 kb) and stronger correlation at large distances (Fig. 3E). This pattern is consistent with the hypothesis that large-scale DSB domains, like hotspots, reflect features of the underlying chromosome architecture that are selectively constrained (13). Furthermore, such large-scale domains presumably reflect operation of factors that work in cis but at a distance from DSB hotspots, i.e., too far to be frequently included in gene conversion tracts and thus not subject to loss through biased gene conversion. Such factors are not expected to evolve as rapidly as hotspots (3, 5, 14, 37).

**Fig. 3.**
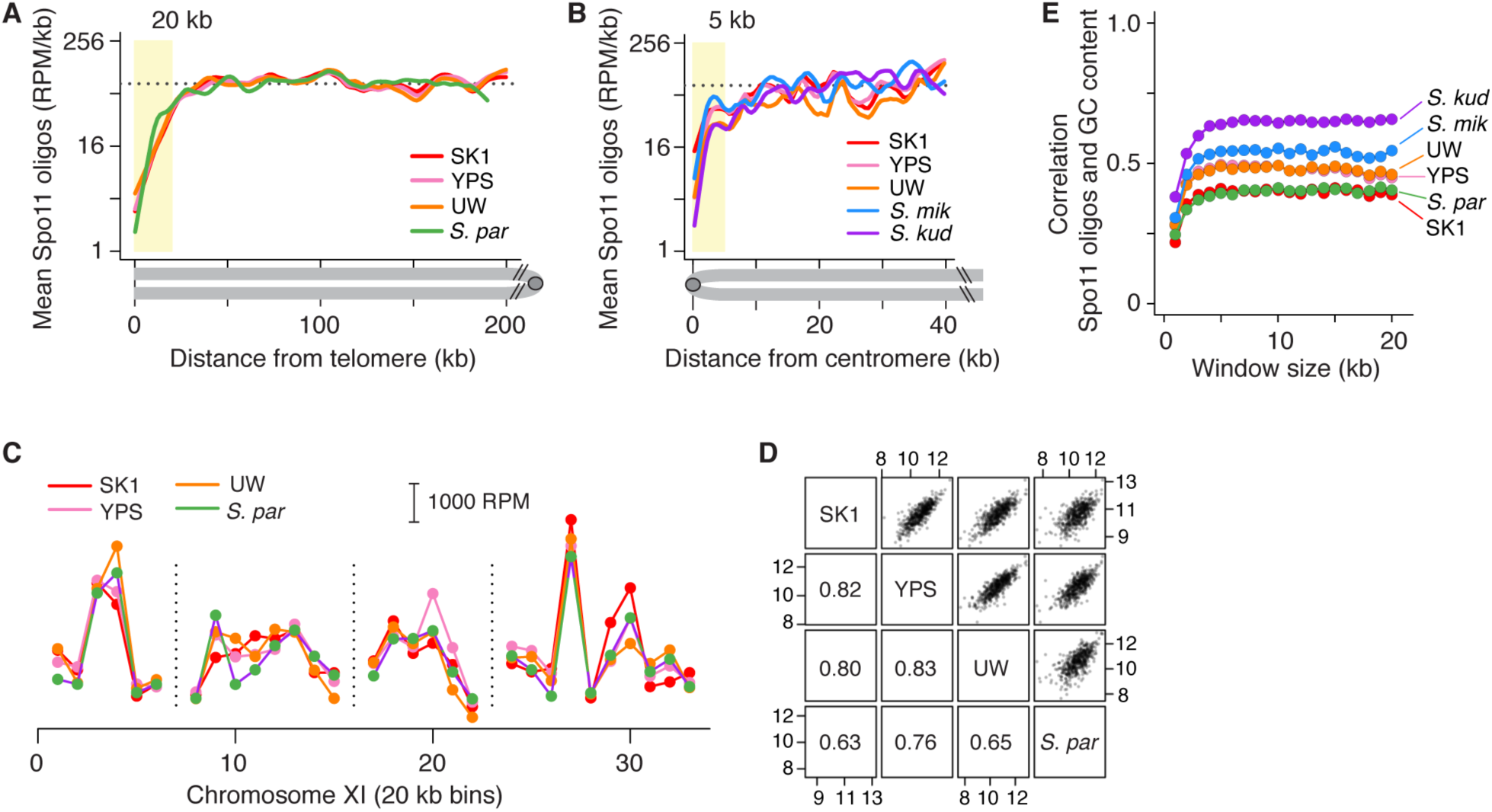
Conservation of large-scale features of the DSB landscape. (A and B) Telomere-proximal and pericentric DSB suppression. Lines are smoothed fit (Lowess) of Spo11-oligo densities in 500 bp bins averaged across 32 chromosome arms. Dashed line, genome average in SK1; yellow shading, DSB suppression zones. Genome assemblies are not complete enough to evaluate telomeric zones for *S. mikatae* or *S. kudriavzevii*. (C,D) Large-scale hot and cold interstitial domains are conserved. Interstitial segments (excluding chromosome ends and peri-centromeres) were defined as syntenic between *S. cerevisiae* and *S. paradoxus* if orthologous genes were in the same order in both species. Spo11-oligo counts were then summed in these syntenic segments divided into 20-kb bins (**Table S5**). A representative example is shown in panel C (vertical dashed lines denote breaks in synteny, many of which reflect unresolved annotation discrepancies) and genome-wide scatter plots and correlation coefficients are in panel D. See also **fig. S5**. (E) Correlation (Pearson’s *r*) between mean Spo11-oligo counts and GC content binned in windows of varying size.

Smaller chromosomes undergo more crossing over per kb than larger chromosomes (38). This behavior can be accounted for by an anticorrelation of DSB density with size (13), which is a patterning effect of a negative feedback circuit that inhibits DSB formation when homologous chromosomes successfully engage one another (39). We find that the negative correlation between Spo11-oligo density and chromosome length is conserved (Fig. 4A, fig. S6). Chromosome bisection and fusion experiments demonstrated that variation in crossover density is determined by chromosome length (38), but this has not been formally tested for DSBs. *S. mikatae* provides a natural experiment: reciprocal translocations between ancestral chromosomes VI, VII, and XVI gave rise to a longer chromosome VI and a shorter chromosome VII in *S. mikatae* than in other *Saccharomyces* species (40) (Fig. 4B). Spo11-oligo density within segments syntenic with the left and right arms of ancestral chromosome VI adopted the Spo11- oligo density predicted by their chromosome length context: density was higher when the segments resided on the short chromosome VI in *S. cerevisiae* but lower when on longer chromosomes in *S. mikatae* (Fig. 4B). Control syntenic segments on similar-length chromosomes exhibited matched Spo11-oligo densities (Fig. 4C). We conclude that whole-chromosome variation in DSB density is a direct consequence of chromosome size per se and is thus extrinsic to the DNA sequence.

**Fig. 4.**
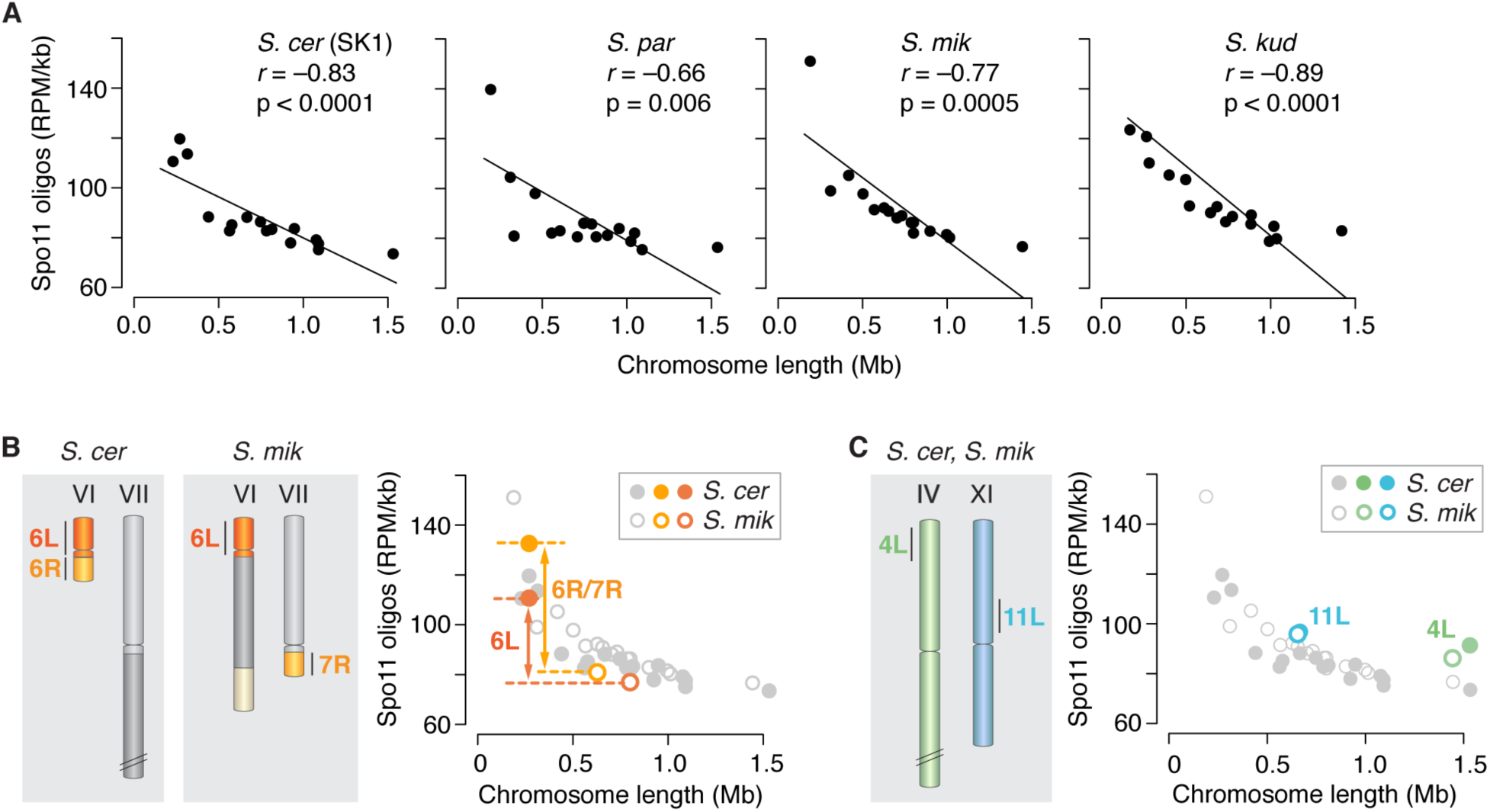
DSB density is influenced by chromosome length. (A) The negative correlation between chromosome length and DSB density is conserved. Each point is one chromosome. (B) A natural experiment demonstrating chromosome length-dependent DSB control. The schematic illustrates a pair of syntenic segments on chromosomes of different size in *S. cerevisiae* and *S. mikatae*. The plot shows that Spo11-oligo density is higher on these segments in *S. cerevisiae* (when on a short chromosome) than in *S. mikatae* (when on longer chromosomes). (C) Control syntenic regions that reside on similarly sized chromosomes have equivalent Spo11-oligo densities in both species.

That yeast hotspot positions are conserved fits the hypothesis that hotspots tend to be stable if Spo11 targets functional genomic elements that are evolutionarily constrained (9). *Drosophila* provides an interesting contrast to reinforce this conclusion: this genus lacks a PRDM9-like system but also does not preferentially target recombination to promoters or known functional elements (in fact, it appears to lack hotspots altogether); accordingly, the fine-scale recombination pattern appears to evolve rapidly (41). A corollary is that evolutionary stability may imply that DSB hotspots have a constrained function(s), even if that function is presently obscure. Interestingly, DSB hotspots are well conserved between the *Schizosaccharomyces* species *S. pombe* and *S. kambucha* (∼0.5% sequence divergence) (42) despite mapping to sites without known function (i.e., in large intergenic regions but not gene promoters (43)).

Strong conservation in Saccharomycetes of relative DSB frequencies among hotspots, across subchromosomal domains, and even across whole chromosomes supports the hypothesis that this conservation traces back to the DSB landscape being shaped by selectively constrained chromosomal features that work combinatorially, hierarchically, and over multiple size scales (13). For example, transcription, replication, telomere and centromere function, sister chromatid cohesion, and chromosome compaction rely on and shape chromosome structures over scales ranging from tens to millions of base pairs. These structures in turn also mold the DSB landscape, so selective pressure to maintain them for gene expression, cell division, and other processes imposes a tendency to conserve the DSB landscape. However, the remarkable strength of conservation across millions of years of evolution leads us to speculate that the specific shape of the yeast DSB landscape may confer fitness benefits. It has long been understood that the recombination distribution is a heritable trait subject to selection (14, 44). We speculate that selective pressures may operate more directly on the DSB landscape genome-wide, perhaps related to accurate meiotic chromosome segregation and/or beneficial effects of disrupting or maintaining linkage groups at various size scales (44, 45).

Finally, we note that available evidence in plants, birds, and dogs — all of which appear to lack a PRDM9-like hotspot targeting mechanism — point to Spo11 acting preferentially at promoters, CpG islands, and/or other genomic elements that are under selective constraint to maintain functions separate from being Spo11 targets (46-48). In finches, high-resolution recombination maps inferred from population genetic data reveal evolutionary stability of recombination hotspots, analogous to *Saccharomyces* spp. but wholly unlike PRDM9-reliant apes or mice (48). Thus, not only is it untrue that recombination initiation landscapes inevitably evolve rapidly, but conservation is likely to be a common pattern for many sexual species.

## Acknowledgments

Submission of Spo11-oligo and MNase-seq data to the Gene Expression Omnibus (GEO) repository is pending. We are grateful to Duncan Greig, Chris Hittinger, Gianni Liti, Ed Louis, Carolin Müller, and Jeremy Roop for providing strains and advice on culturing *Saccharomyces* species, Agnes Viale and the Integrated Genomics Operation (MSKCC) for sequencing, Nick Socci (Bioinformatics Core Facility, MSKCC) for mapping Spo11-oligo reads, Stewart Shuman for gifts of T4 RNA ligase, and Sarah Kim, Julian Lange, Neeman Mohibullah, Sam Tischfield, and Xuan Zhu for helpful discussions on the bioinformatic analyses. We thank Molly Przeworski for discussions and communicating data prior to publication, and for providing helpful comments on the manuscript. This work was supported by NIH grant R01 GM058673. IL was supported in part by NIH fellowship F31 GM097861. SK is an investigator of the Howard Hughes Medical Institute.

